# Evaluating the tea bag method as a potential tool for detecting the effects of added nutrients and their interactions with climate on litter decomposition

**DOI:** 10.1101/2021.01.28.428520

**Authors:** Taiki Mori, Toru Hashimoto, Yoshimi Sakai

## Abstract

It is acknowledged that exogenous nutrient addition often stimulates early-stage litter decomposition in forests and late-stage decomposition is generally suppressed by nitrogen addition, whereas the interactive effects of nutrient addition and abiotic environmental factors, such as climate, on decomposition remain unclear. The tea bag method, which was developed to provide the decomposition rate constant *k* of early-stage decomposition and stabilization factor *S* of labile materials in the late stage, is a potentially useful tool for examining the impacts of nutrient addition on both early- and late-stage litter decomposition and their interactions with climate. At a long-term (38-year) continuous fertilization experimental site (an *Abies sachalinensis* Fr. Schmidt stand) in Hokkaido, Japan, we examined whether a standard tea bag method protocol was sufficiently sensitive to reveal any impacts of nutrient addition on early- and late-stage decomposition. In addition, we tested the interactive effects of nutrient addition and climate on litter decomposition. The short incubation period of the tea bag method (ca. 90 days) enabled us to obtain decomposition data from the same location at three different times in a year, i.e., early summer, midsummer, and winter, providing an opportunity to test interactive effects. We demonstrated that the decomposition rate of rooibos tea and the decomposition rate constant *k* of early-stage decomposition were clearly stimulated by fertilization in midsummer, but no impacts were detected in other seasons, probably because the relative importance of nutrient availability was elevated in midsummer, during which decomposition rates were less constrained by temperature and moisture. The green tea decomposition rate and stabilization factor *S*, an index related to late-stage decomposition, were unaffected by fertilization. This was probably because the tea bag method does not take into account lignin degradation, which is considered a key factor controlling late-stage litter decomposition. Overall, the present study (i) successfully determined the interactive effects of nutrient addition and climate factors on litter decomposition by making full use of the tea bag method, and (ii) the results suggest that the tea bag method can be a suitable tool for examining the direct effects of nutrient addition and their interactions with environmental factors on early-stage litter decomposition, but not those on late-stage decomposition.

## Introduction

Since the industrial revolution, anthropogenic nutrient inputs into ecosystems have been elevated substantially, which could have large impacts on ecosystem function (Galloway et al., 2008). Litter decomposition is an essential process that controls both nutrient recycling and carbon (C) dynamics in forest ecosystems. Litter decomposition can be divided into two stages—the early stage, when soluble compounds and non-lignified cellulose and hemicellulose are degraded, and the late stage, during which lignified tissues are degraded. It is essential to understand how nutrient loading, especially of nitrogen (N) and phosphorus (P), affects both early- and late-stage litter decomposition.

It is widely known that late-stage decomposition is generally inhibited by N addition (Berg, 1986; Janssens et al., 2010; Knorr, Frey, & Curtis, 2005) because of the following possible reasons, which are not necessarily mutually exclusive: (i) N addition causes the production of chemically recalcitrant materials (Berg and Matzner, 1997, but see Rinkes et al., 2016); (ii) N toxicity, high-salt conditions, or acidification caused by N addition negatively affects microbial activity; (iii) added N causes microbes to stop acquiring N from organic matter (Craine, Morrow, & Fierer, 2007); and (iv) microbial community changes can be caused by N addition (Bonner et al., 2019; Ramirez, Craine, & Fierer, 2012). Similarly, P addition may also suppress late-stage litter decomposition (DeForest, 2019; Mori et al., 2015), although these mechanisms are not well understood. Conversely, early-stage litter decomposition (or the decomposition of litter with low lignin content) is often stimulated by the additions of N (Berg & Matzner, 1997; Fog, 1988; Knorr et al., 2005) and P (Mori et al., 2015), probably because the nutrient limitations of decomposers are relieved by nutrient addition. Recent meta-analyses have demonstrated that the activities of enzymes that degrade cellulose and hemicellulose are stimulated by N, whereas N addition suppresses oxidative enzymes that have an important role in degrading lignin (Jian et al., 2016; Xiao, Chen, Jing, & Zhu, 2018), successfully explaining the contrasting impacts of N loading on the early- and late-stages of litter decomposition.

Thus, it is acknowledged that exogenous nutrient additions often stimulate early-stage litter decomposition in forests, whereas late-stage decomposition is generally suppressed by N addition. However, the interactive effects of nutrient addition and various abiotic environmental factors, especially climatic factors such as temperature and moisture, on decomposition remain unclear. For example, the relative importance of nutrient availability may be elevated under hotter and wetter conditions, during which decomposition is less restricted by temperature and moisture. If this is the case, the stimulating effects of nutrient addition on early-stage litter decomposition may be more intensive under hotter and wetter conditions. The investigation of these effects requires data sets of litter decomposition rates from fertilization studies conducted using standardized materials under different environmental conditions, because the quality of litter significantly affects the response of decomposition to added nutrients (Knorr et al., 2005), thus masking the impacts of nutrient addition and environmental factors.

Tea bag decomposition is a cost-effective method using standardized materials that can be employed to obtain indices related to both early- and late-stage decomposition with a single measurement after a 90-day field incubation. Using this method, the limit value (i.e., the point at which the decomposition process either continues at a very low rate or possibly stops; Berg et al., 2010) of the hydrolysable fraction of teas can be expressed as the mass-loss ratio of green tea, assuming that (i) the decay rate of the hydrolysable fraction of green tea becomes nearly zero at 90 days and (ii) the undecomposed hydrolysable fraction at 90 days is stabilized and remains undecomposed over the long term. The ratio of the undecomposed fraction to the total hydrolysable fraction is defined as the “stabilization factor *S*,” which is considered to indicate long-term carbon storage (Fujii et al., 2017). On the other hand, the decomposition constant *k* of early-stage decomposition is determined by calculating the decomposition rate of rooibos tea using an asymptotic model, assuming that the stabilization factor *S* of rooibos tea is the same as that of green tea (for more details, see the Materials and Methods section). This method has been proposed as an effective approach for collecting comparable globally distributed data (Keuskamp, Dingemans, Lehtinen, Sarneel, & Hefting, 2013), and is used by an increasing number of researchers (Djukic et al., 2018; Fujii et al., 2017; Mueller et al., 2018; Petraglia et al., 2019; Suzuki et al., 2019).

The tea bag method constitutes a potentially useful tool for examining the direct effects of nutrient additions on both the early- and late-stages of litter decomposition, requiring less effort than other methods. In addition, it can also enable researchers to determine the interactions between nutrient additions and various environmental factors with respect to the decomposition of litter, owing to the well-standardized materials used in the method. In the present study, we investigated whether the standard protocol of the tea bag method was sufficiently sensitive to reveal (i) any impacts of nutrient addition on indices related to early- (i.e., decomposition constant *k*) and late-stage (i.e., stabilization factor *S*) decomposition and (ii) any interactive effects of nutrient addition and abiotic environmental factors on decomposition. To focus on the interactive effects of nutrient addition and climate factors, which are the most important abiotic environmental factors controlling organic matter decomposition, we tried to minimize site effects other than climate by performing decomposition experiments at the same study site but in different seasons of the year, i.e., early summer, midsummer, and winter. The short incubation term (ca. 90 days) of the protocol made this approach possible.

We hypothesized that early-stage decomposition rates (indicated by the mass loss ratio of rooibos tea and decomposition constant *k*) are stimulated by nutrient addition, which relieves the nutrient limitation of soil microbes. On the other hand, the stabilization factor *S*, an index related to late-stage decomposition, as well as the mass loss ratio of green tea, should be unaffected by nutrient addition because the tea bag method and the tea bag indices do not consider lignin degradation, the most important fraction controlling late-stage decomposition and its response to N addition. We also hypothesized that the impact of nutrient addition on early-stage litter decomposition would be more remarkable in hotter and wetter climates.

## Materials and methods

### Study site

The present study was performed in a portion of a long-term fertilization experimental site in the experimental forest of the Hokkaido Research Center, Forestry and Forest Products Research Institute, Sapporo, Japan (42°59′N, 141°23′E) (Aizawa et al., 2012; Furusawa, Nagakura, Aizawa, & Ito, 2019). The mean annual temperature and annual precipitation at the study site in 2015, as determined using Agro-Meteorological Grid Square Data, NARO, were 8.5 °C and 1071 mm, respectively (Fig. S1). An *Abies sachalinensis Fr. Schmidt* (Sakhalin fir) stand was established at the site in 1973, with four sub-plots: C1 (control 1), C2 (control 2), NP (fertilized with N and P), and NP6Y (fertilized with N and P only for the first 6 years). Fertilization with N and P was initiated in 1978, using ammonium sulfate and lime superphosphate. The average amounts of added N and P were 114 kg N ha^−1^ year^−1^ and 33 kg P ha^−1^ year^−1^ for the first 6 years, and 122 kg N ha^−1^ year^−1^ and 34 kg P ha^−1^ year^−1^ over the whole experimental period (38 years). Impacts of this fertilization on diameter at breast height, tree height, soil pH, and soil total C content have been reported by a previous study (Aizawa et al., 2012) (Fig. 1).

**Fig. 1.**
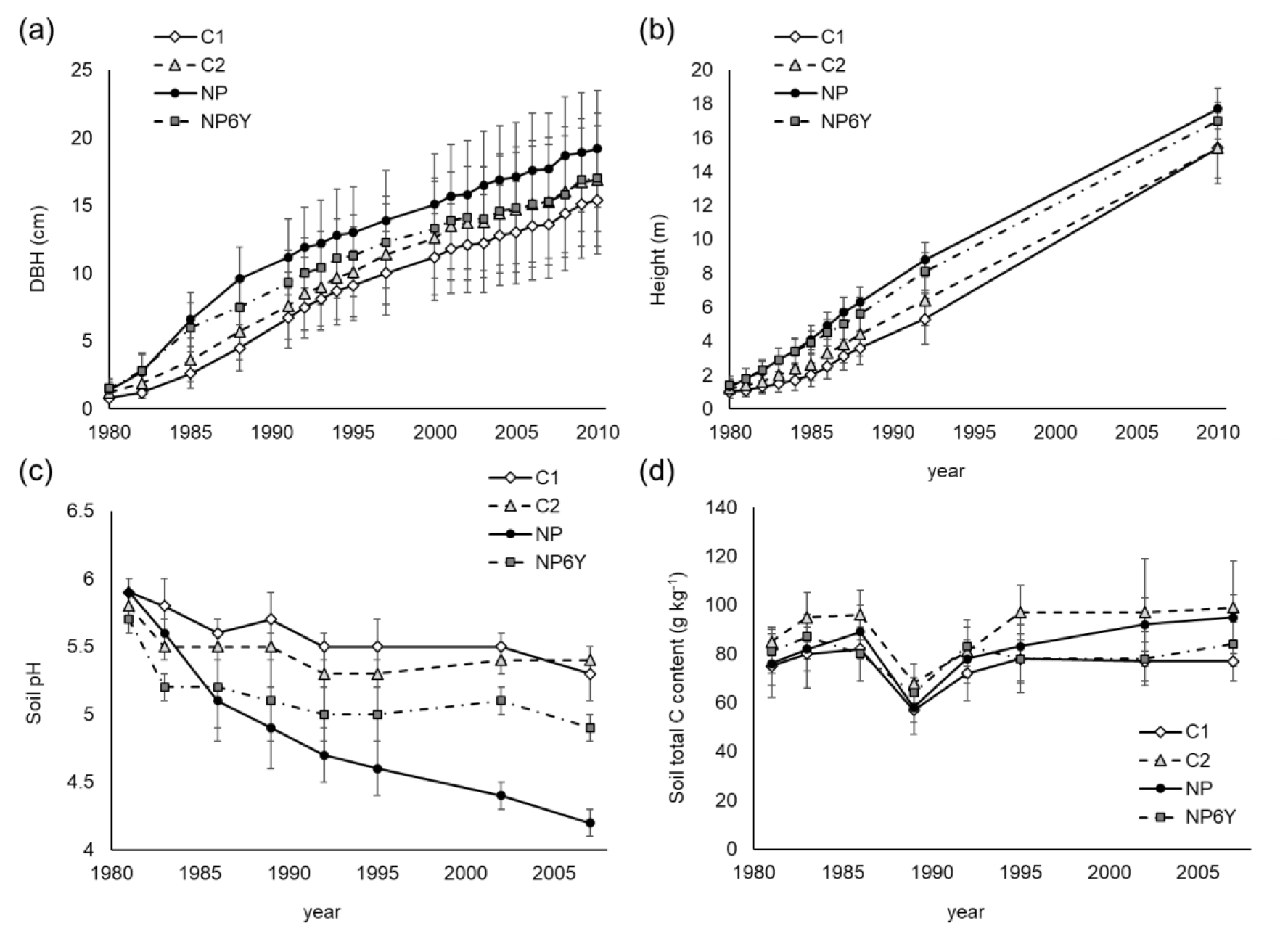
Changes in (a) diameter at breast height (DBH), (b) tree height, (c) soil pH, and (d) soil total C content after N and P fertilization at the study sites. Data were taken from Aizawa et al. (2012). Error bars indicate standard deviations. C1, control 1; C2, control 2; NP, fertilized with N and P; NP6Y, fertilized with N and P only for the first 6 years.

### Tea bag incubation

Tea bag incubations were performed in early summer (91 days, from 1 June to 31 August), midsummer (90 days, from 2 July to 30 September), and winter (89 days, from 5 November to 2 February). Cumulative precipitations and accumulated temperatures higher than zero during the field incubation experiments were calculated using data from Agro-Meteorological Grid Square Data, NARO (Fig. 2). Basically, we followed the protocol from Keuskamp et al. (2013). Green (EAN: 87 22,700 05552 5) and rooibos (EAN: 87 22,700 18,843 8) teabags produced by Lipton (0.25-mm mesh) were used for the experiment. We evenly divided each sub-plot into four areas, in each of which two replicate pairs of tea bags (green and rooibos tea bags) were buried in the soil at 8-cm depth. Tea bags were retrieved approximately 90 days later. The bags were oven dried at 70 °C for 48 h, and dry weights were determined.

**Fig. 2.**
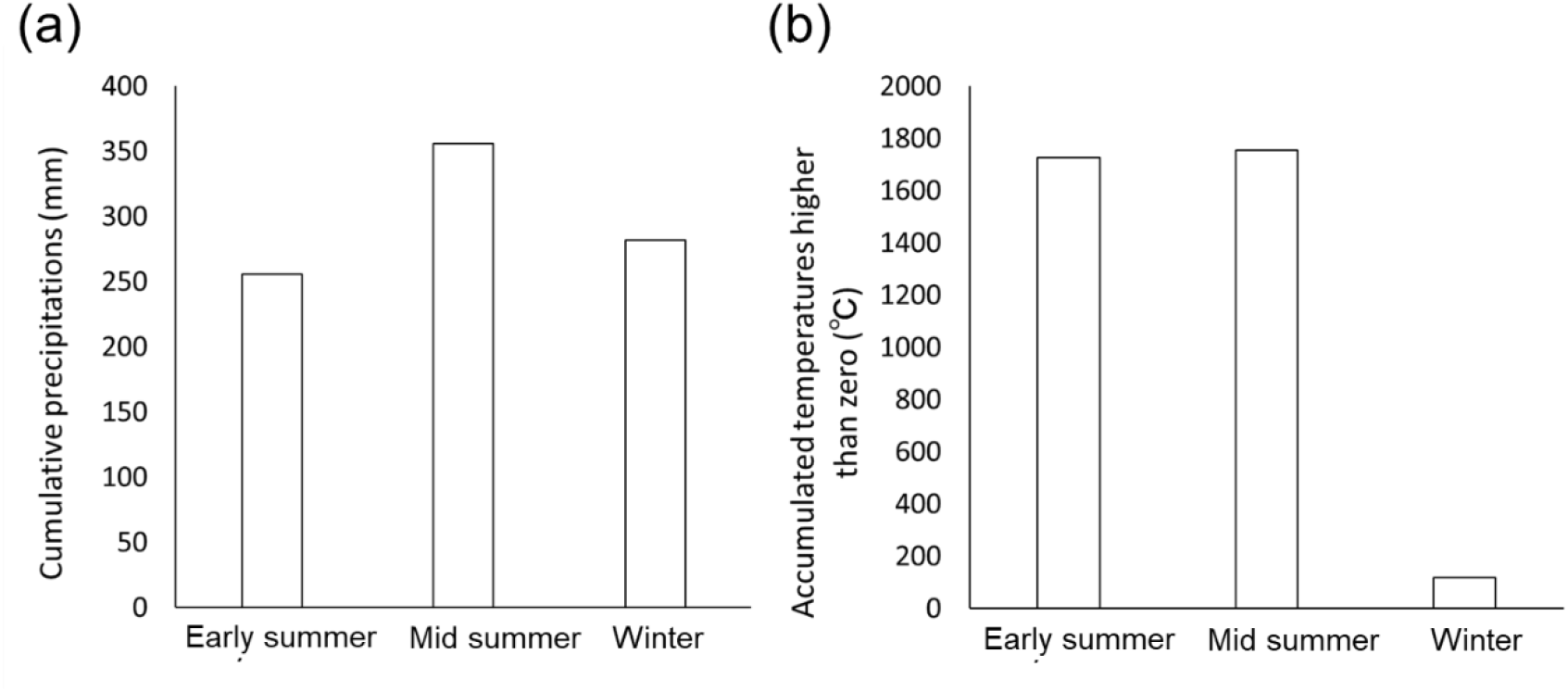
Climate data during the experiment. (a) Cumulative precipitation and (b) accumulated temperatures higher than zero during the field incubation experiments were calculated using data from Agro-Meteorological Grid Square Data, NARO.

### Calculation of tea bag indices

Following Keuskamp et al. (2013), we calculated the tea bag indices, i.e., the stabilization factor *S* and decomposition constant *k*. The stabilization factor *S* was calculated as the relative amount of labile fractions of green tea that were stabilized and transformed to recalcitrant fractions during decomposition (Keuskamp et al., 2013). *S* can be calculated as follows, assuming that (i) the decomposition of the labile fractions of green tea was complete during the incubation period and (ii) the remaining fractions were transformed into recalcitrant fractions:

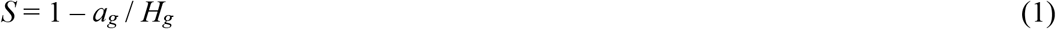

where *a_g_* is the decomposed fraction and *H_g_* (0.842; Keuskamp et al. 2013) is the hydrolysable fraction of green tea. The decomposed fraction of green tea was determined after incubating green tea, as follows:

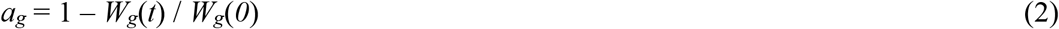

where *W_g_*(*t*) is the weight of the green tea after incubation time *t* and *W_g_*(*0*) is the weight of the green tea before incubation.

The decomposition process of organic matter, including both easily degradable and recalcitrant compounds, can be approximated as follows (Wieder & Lang, 1982):

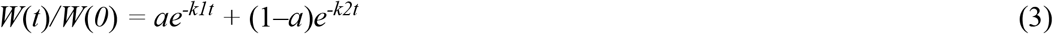

where *W*(*t*) is the weight of the substrate after incubation time *t*, *W*(*0*) is the weight of the substrate before incubation, *a* and (*1-a*) are the labile and recalcitrant fractions of the litter, respectively, and *k_1_* and *k_2_* are the decomposition rate constants of the labile and recalcitrant fractions, respectively. Because the weight loss of the recalcitrant fraction is generally negligible during short-term field incubations, Eq. (3) can be reduced as follows, assuming *k_2_* = 0:

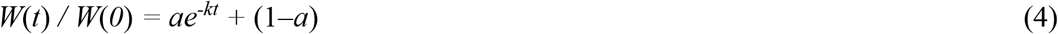

For the tea bag indices, the decomposition rate of rooibos tea is used to calculate decomposition constant *k*. The decomposable fraction of rooibos tea (*a_r_*) can be calculated using the value of *S*, assuming that the same ratio of labile fractions is stabilized in rooibos tea as in green tea:

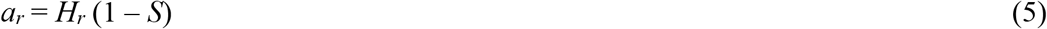

where *H_r_* is the hydrolysable fraction of rooibos tea (0.552; Keuskamp et al. 2013). The decomposition constant *k* is calculated by applying *W*_r_(0), *W_r_*(*t*), *t*, and *a_r_* to the exponential decay function given in Eq. (4) as follows:

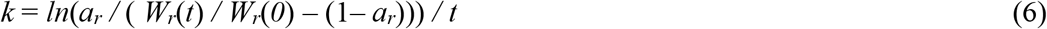

where *W_r_*(*t*) is the weight of the rooibos tea after incubation time *t* and *W_r_*(*0*) is the weight of the rooibos tea before incubation.

### Statistics

Two-way ANOVA followed by Tukey’s *post hoc* tests were used to determine the significance of the impacts of fertilization and season, assuming normal distribution of the data. If any interaction was significant, simple main effect analyses (Tukey’s *ad-hoc* tests) were performed to detect the factor(s) causing the interaction.

The correlation between decomposition constant *k* and stabilization factor *S* was tested using Pearson’s correlation test. All eight (six for winter) samples were used in the analysis. The correlation analysis was carefully done; we compared the results of the correlation analysis with correlations of simulated *k* and *S* that were calculated from random data generated from real data obtained from the field experiment. This comparison was needed because *S*, the function of green tea decomposition rate, and *k*, the function of the decomposition rates of both green and rooibos teas, could be correlated even if the data for tea bag decomposition rates were randomly distributed. The simulation data were generated using the average and standard deviation of the field data, assuming a standard distribution. Data obtained from midsummer were used. Each generated simulation datum of the ratio of green tea mass remaining was paired up with a datum for rooibos tea. The generated pairs were not correlated with each other (see Fig. S3). Using this simulation data, the decomposition constant *k* and stabilization factor *S* were calculated. All statistical analyses were performed using R software (R Core Team, 2019).

## Results

The decomposition rates of the green and rooibos teas were within the range of data taken from other study sites in Japan (Fig. S2), although the decomposition rates of both green and rooibos teas in winter might have been slightly elevated at our study site (Fig. S2a, b).

Green tea decomposition was significantly affected by the season but not by fertilization (two-way ANOVA, Fig. 3a). Tukey’s multi-comparison tests showed that the decomposition of green tea was quicker in midsummer than in early summer or winter. Season and fertilization influenced the decomposition of rooibos tea with a significant interaction (two-way ANOVA, Fig. 3b). The simple main effect analysis suggested that (i) the effect of fertilization on rooibos tea decomposition was significant only in midsummer and (ii) decomposition rates were slower in winter compared with early and mid-summer. Although the incubation periods were slightly different among seasons (91, 90, and 89 days in early summer, midsummer, and winter, respectively), statistical analyses using the normalized data (mass loss ratio per day) demonstrated that the results were not affected by the differences in incubation periods.

**Fig. 3.**
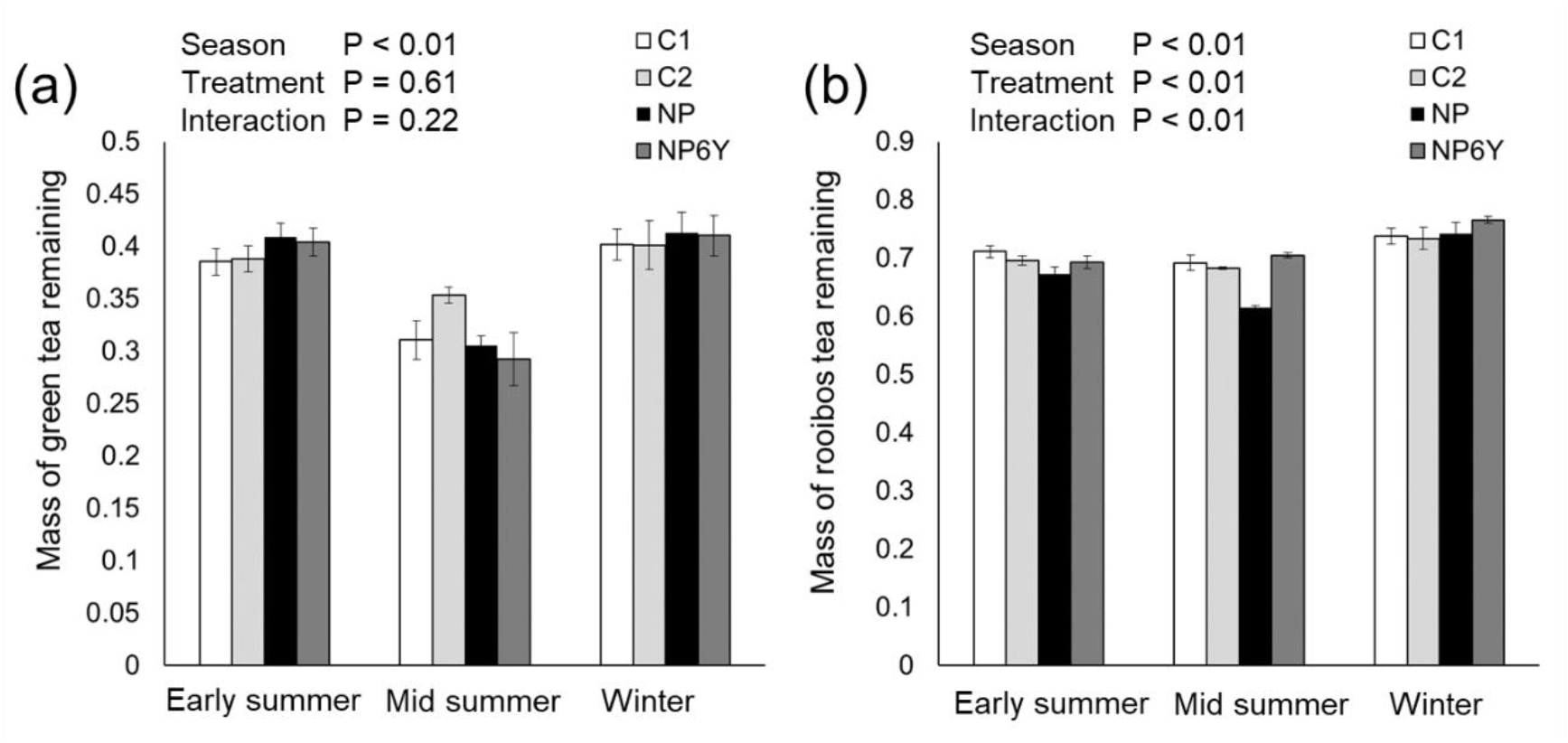
Effects of season and fertilization on the mass of (a) green tea and (b) rooibos tea remaining after incubation. Each error bar indicates the standard error of four replicates.

The impacts of fertilization and season on stabilization factor *S* were the same as those on green tea decomposition (Fig. 4a), because *S* was calculated using the mass loss ratio of green tea multiplied by a constant factor (see Eq. 1 and 2). Meanwhile, the impacts of fertilization and season on decomposition constant *k* are affected by the decomposition rates of both green and rooibos teas (see Eq. 4 and 5). The decomposition constant *k* was affected by an interaction between fertilization and season (two-way ANOVA, Fig. 4b). A simple main effect analysis suggested that the effects of fertilization on rooibos tea decomposition were significant only in midsummer.

**Fig. 4.**
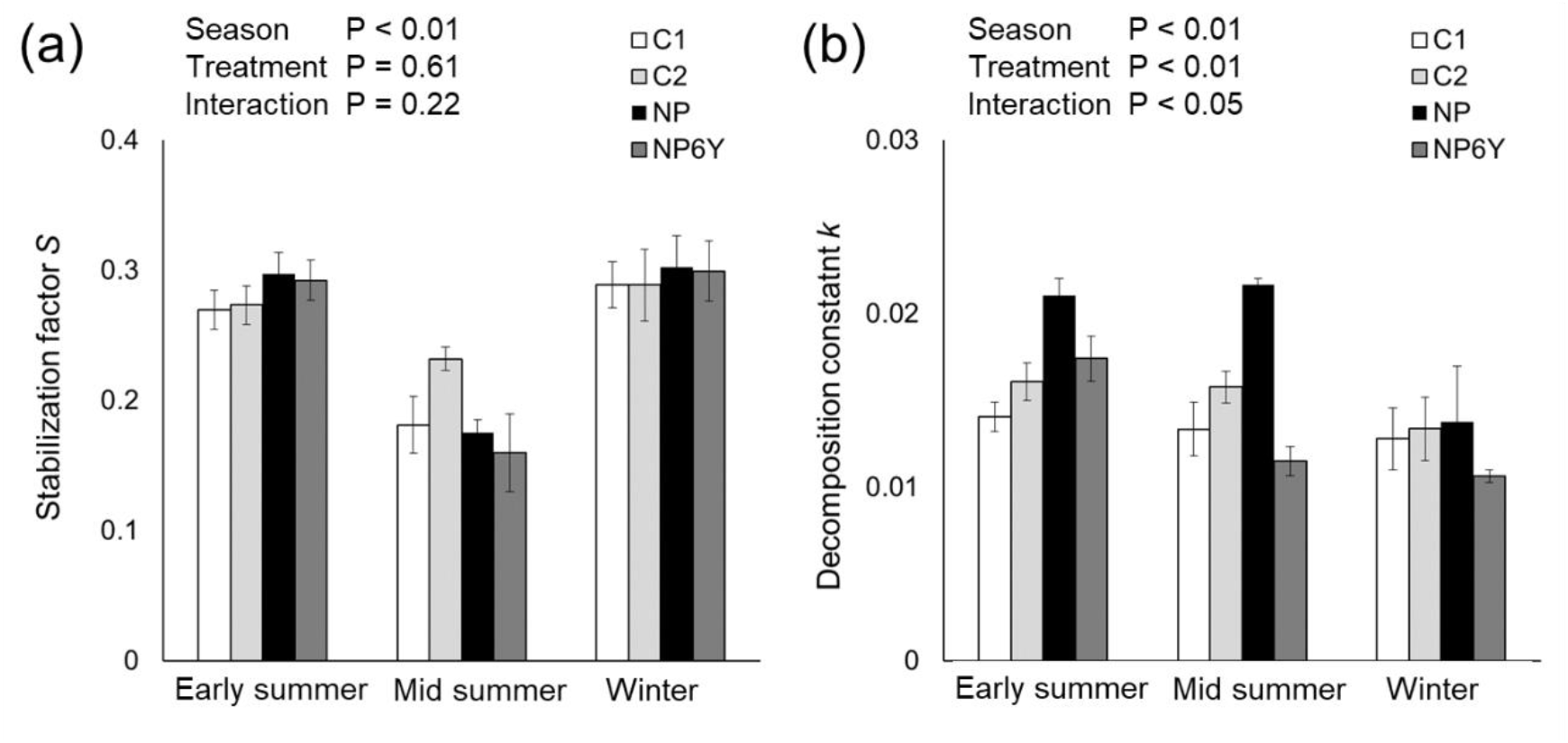
Effects of season and fertilization on (a) stabilization factor *S* and (b) decomposition constant *k*. Each error bar indicates the standard error of four replicates.

We found significant positive correlations between *k* and *S* in all four treatments in midsummer (Fig. 5b) and in the NP plot in winter (Fig. 5c). The simulation also demonstrated a similar pattern—i.e., a significant positive correlation between simulated *k* and *S* (Fig. 5d).

**Fig. 5.**
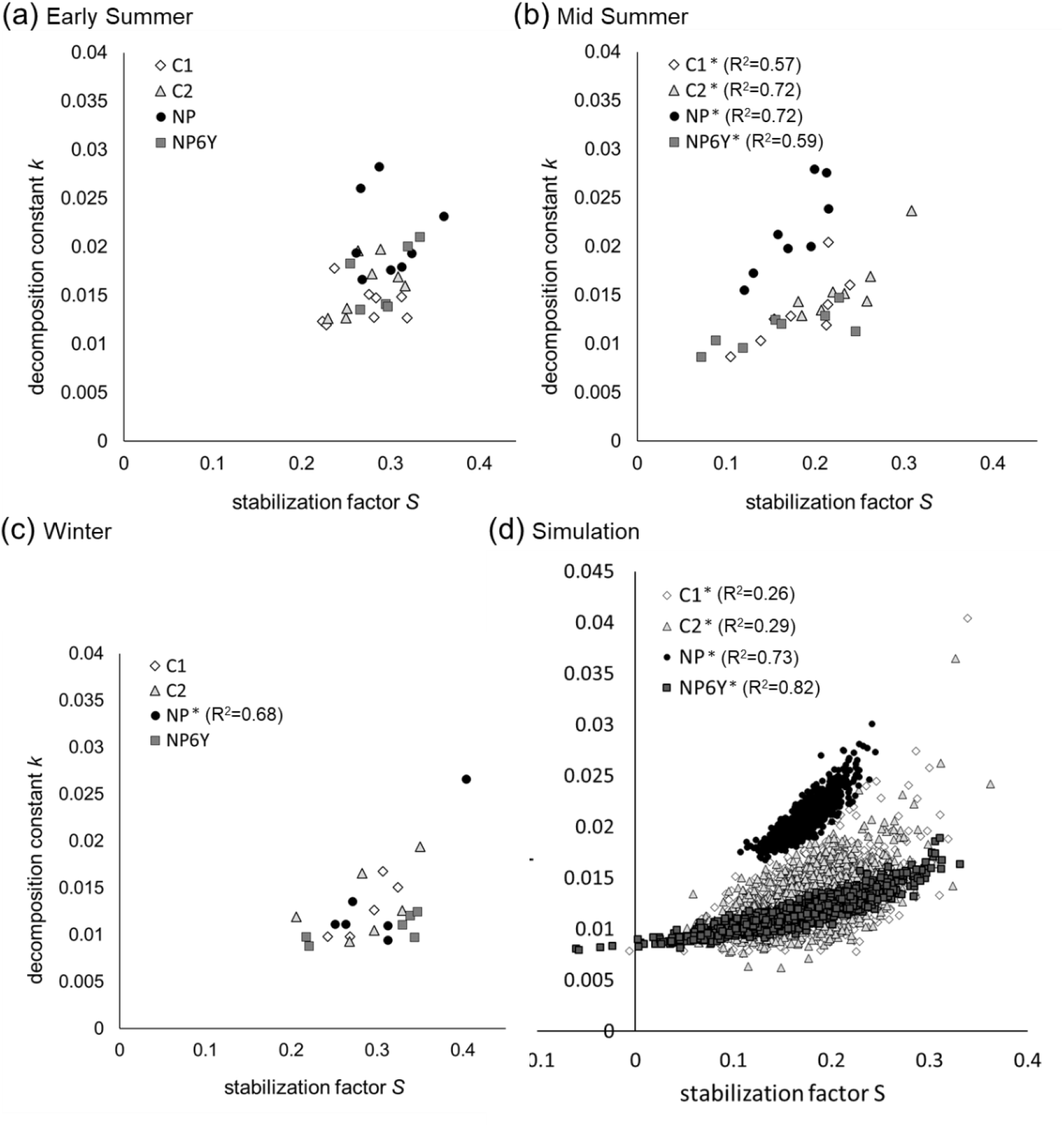
Relationships between decomposition constant *k* and stabilization factor *S* in (a) early summer, (b) midsummer, and (c) winter and (d) using simulated data. Asterisks (*) indicate statistically significant correlations (P<0.05, Pearson’s correlation). Simulation data (n =1000) of ratios of green tea and rooibos tea mass remaining were generated using averages and standard deviations of field data obtained in late summer, assuming a standard distribution. Each datum of the ratio of green tea mass remaining was paired with a datum of rooibos tea. We confirmed that the generated pairs did not exhibit a correlation (see Fig. S2). Using the data, decomposition constant *k* and stabilization factor *S* were calculated. The generated *k* and *S* values exhibited significant positive correlations (d), indicating that positive correlations can be automatically obtained.

## Discussion

### Impacts of nutrient addition on early-stage litter decomposition and interactions with climate

As we hypothesized, nutrient addition significantly stimulated early-stage litter decomposition in midsummer, as indicated by the elevated mass loss ratio of rooibos tea and decomposition constant *k*. This result is consistent with earlier studies, which reported that litters with higher nutrient contents, or those amended with nutrients, had higher decomposition rates (Berg & Matzner, 1997; Carreiro, Sinsabaugh, Repert, & Parkhurst, 2000; Fog, 1988; Mori et al., 2015), probably because microbial nutrient shortages were relieved. The fact that rooibos tea decomposition was stimulated only in midsummer but not in early summer or winter (Fig. 2b) indicated that the tea bag method successfully detected the interactive effects of nutrient addition and climate on litter decomposition. The relative importance of nutrient availability was probably elevated in midsummer, during which decomposition rates were less restricted by temperature than during winter or by higher moisture than in early summer (Fig. 2). This suggests that the impacts of fertilization may be stronger in hotter and wetter climate zones, such as tropical rain forests. Thus, we successfully demonstrated that the tea bag method is a suitable tool for examining the direct effects of nutrient addition and their interactions with environmental factors on early-stage litter decomposition. Larger amounts of data from fertilization studies using tea bag methods conducted at various sites would enable the clarification of the interactive effects of fertilization and environmental factors other than climate, such as soil and background vegetation types.

### Impacts of nutrient addition on late-stage litter decomposition

As we hypothesized, green tea decomposition rates and the stabilization factor *S* were not influenced by experimental nutrient addition. The lack of response of decomposition rates to nutrient addition does not necessarily indicate a methodological unsuitability, but in addition to observations from the field experiment, we have a theoretical basis for assuming that the tea bag method has unsuitable characteristics for determining the impacts of nutrient loading on late-stage litter decomposition. The tea bag method focuses on the hydrolysable fraction, and it is assumed that lignin degradation is negligible during the 90-day incubation of the teas (Keuskamp et al., 2013). As a result, lignin degradation is not considered when determining stabilization factor *S*. Thus, the response of lignin decomposition to nutrient addition is undetectable using the tea bag method. Because it is well acknowledged that lignin, or the unhydrolysable fraction of litter, is a key factor in controlling late-stage litter decomposition, and N addition suppresses the degradation of lignin (Jian et al., 2016; Rinkes et al., 2016), the tea bag method would be unsuitable for assessing the impacts of nutrient addition on long-term C storage, although this method may be a useful tool for estimating C stability.

Our study demonstrated that the decomposition rates of green tea were significantly influenced by season, with quicker decomposition in midsummer than in other seasons (Fig. 3a). Accordingly, the stabilization factor *S* was smaller in midsummer than in other seasons (Fig. 4a). This was attributed to lower temperatures in winter (Fig. 2b) and smaller amounts of precipitation in early summer (Fig. 2a) compared with midsummer. According to the tea bag method, this result can be interpreted as larger amounts of labile fractions being transformed into recalcitrant fractions (Keuskamp et al., 2013) in early summer and winter compared to midsummer. However, it is possible that the decomposition of labile fractions of green tea was retarded by less favorable conditions for decomposers, and thus did not reach its limit value in early summer and winter. The results of several previous studies may support this, as higher decomposition rates of green tea were reported on the 150^th^ day compared to the 90^th^ day (Wang et al., 2019) as well as on the 136^th^ day compared to the 89^th^ day (Seelen et al., 2019), indicating that green tea decomposition did not reach its limit values at 90 days. If this was the case, *S* was overestimated in winter and early summer (Fig. S4), and hence, the decomposition rate *k* was also overestimated (Fig. S4). This idea should be tested in future studies.

### Raw decomposition data vs k for evaluating the impacts of nutrient addition

Although the direct impacts of nutrient addition and its interaction with climate factors were successfully detected using both raw decomposition data and the decomposition rate constant *k*, we have several reasons to consider that the raw decomposition data from the tea bag experiment may be better for evaluating the impacts of nutrient addition on litter decomposition. As discussed earlier, *S* might have been overestimated because the decomposition of the hydrolysable fraction might not have reached its limit value, in which case, *k* would also be overestimated (Fig. S4). In addition, a premise for calculating *k*, that the stabilization factor is the same for both green and rooibos teas, was not verified, which also risks the over- or under-estimation of *k*. Furthermore, plotting the relationship between *k* and *S*, as several previous studies have reported (Becker & Kuzyakov, 2018; Keuskamp et al., 2013; Macdonald et al., 2018; Seelen et al., 2019), may be misleading. In the present study, we found significant positive correlations between *k* and *S* in all four treatments in midsummer and in the NP plot in winter (Fig. 5bc). There is a risk of misinterpreting positive correlations according to the definitions of the tea bag indices, because a positive correlation was also obtained using randomly generated data (Fig. 5d). Although these concerns regarding over- and under-estimation or misinterpretation are not always valid, it may be true that calculating *k* values to evaluate the impacts of nutrient addition on litter decomposition rates needs great care for the results to be adequately interpreted. We suggest that raw rooibos tea decomposition rates may be a better indicator when assessing the impacts of nutrient addition on litter decomposition.

## Conclusion

Using the tea bag method, we successfully demonstrated that nutrient addition and climate factors have an interactive effect on litter decomposition. We also suggest that the tea bag method may be a suitable tool for examining the direct effects of nutrient addition and its interactions with environmental factors on early-stage organic matter decomposition, but not those on late-stage decomposition.

## Acknowledgement

This research was funded by JSPS KAKENHI Grant Number JP19K15879.

## Notes

Conflict of Interest: None declared.

### Competing Interest Statement

The authors have declared no competing interest.

## References

Aizawa, S., Ito, E., Hashimoto, T., Sakata, T., Sakai, H., Tanaka, N.,… Sanada, M. (2012). Influence of fertilization on growth of 37-years-old plantations of Abies sachalinensis, Picea jezoensis, Picea glehnii and Betula maximowicziana (in Japanese). Boreal For Res, 60, 93–99.

Becker, J. N., & Kuzyakov, Y. (2018). Teatime on Mount Kilimanjaro: Assessing climate and land-use effects on litter decomposition and stabilization using the Tea Bag Index. Land Degradation and Development, 29(8), 2321–2329. doi:10.1002/ldr.2982

Berg, B., Davey, M. P., de Marco, A., Emmett, B., Faituri, M., Hobbie, S. E.,… Virzo De Santo, A. (2010). Factors influencing limit values for pine needle litter decomposition: A synthesis for boreal and temperate pine forest systems. Biogeochemistry, 100(1), 57–73. doi:10.1007/s10533-009-9404-y

Berg, B, & Matzner, E. (1997). Effect of N deposition on decomposition of plant litter and soil organic matter in forest systems. Environ. Rev., 5, Environ. Rev. 5: 1–25 (1997). doi:10.1139/a96-017

Berg, Björn. (1986). Nutrient release from litter and humus in coniferous forest soils: a mini review. Scandinavian Journal of Forest Research, 1(June), 359–369. doi:10.1080/02827588609382428

Bonner, M. T., Castro, D., Schneider, A. N., Sundström, G., Hurry, V., Street, N. R., & Näsholm, T. (2019). Why does nitrogen addition to forest soils inhibit decomposition? Soil Biology and Biochemistry, 117(March), 107570. doi:10.1016/j.soilbio.2019.107570

Carreiro, M. M., Sinsabaugh, R. L., Repert, D. A., & Parkhurst, D. F. (2000). Microbial Enzyme Shifts Explain Litter Decay Responses ToEnzyme, Microbial Explain, Shifts Decay, Litter To, Responses Deposition, Simulated Nitrogen. Ecology, 81(9), 2359–2365.

Craine, J. M., Morrow, C., & Fierer, N. (2007). Microbial nitrogen limitation increases decomposition. Ecology, 88(8), 2105–2113. doi:10.1890/06-1847.1

DeForest, J. L. (2019). Chronic phosphorus enrichment and elevated pH suppresses Quercus spp. leaf litter decomposition in a temperate forest. Soil Biology and Biochemistry, 135(January), 206–212. doi:10.1016/j.soilbio.2019.05.005

Djukic, I., Kepfer-rojas, S., Kappel, I., Steenberg, K., Caliman, A., Paquette, A.,… Schaub, M. (2018). Early stage litter decomposition across biomes, 629, 1369–1394. doi:10.1016/j.scitotenv.2018.01.012

Fog, K. (1988). The effect of added nitrogen on the rate of decomposition of organic matter. Biological Reviews, 63(3), 433–462. doi:10.1111/j.1469-185X.1988.tb00725.x

Fujii, S., Mori, A. S., Koide, D., Makoto, K., Matsuoka, S., Osono, T., & Isbell, F. (2017). Disentangling relationships between plant diversity and decomposition processes under forest restoration. Journal of Applied Ecology, 54(1), 80–90. doi:10.1111/1365-2664.12733

Furusawa, H., Nagakura, J., Aizawa, S., & Ito, E. (2019). Effects of repeated fertilization and liming on soil microbial biomass in Betula maximowicziana Regel and Abies sachalinensis Fr. Schmidt stands in Japan. Landscape and Ecological Engineering, 15(1), 101–111. doi:10.1007/s11355-018-0366-x

Galloway, J. N., Townsend, A. R., Erisman, J. W., Bekunda, M., Cai, Z., Freney, J. R.,… Sutton, M. a. (2008). Transformation of the Nitrogen Cycle: Recent Trends, Questions, and Potential Solutions. Science, 320(5878), 889–892. doi:10.1126/science.1136674

Janssens, I., Dieleman, W., Luyssaert, S., Subke, J., Reichstein, M., Ceulemans, R.,… Law, B. (2010). Reduction of forest soil respiration in response to nitrogen deposition. Nature Geoscience, 3(5), 315–322. doi:10.1038/ngeo844

Jian, S., Li, J., Chen, J., Wang, G., Mayes, M. A., Dzantor, K. E.,… Luo, Y. (2016). Soil extracellular enzyme activities, soil carbon and nitrogen storage under nitrogen fertilization: A meta-analysis. Soil Biology and Biochemistry, 101, 32–43. doi:10.1016/j.soilbio.2016.07.003

Keuskamp, J. A., Dingemans, B. J. J., Lehtinen, T., Sarneel, J. M., & Hefting, M. M. (2013). Tea Bag Index: A novel approach to collect uniform decomposition data across ecosystems. Methods in Ecology and Evolution, 4(11), 1070–1075. doi:10.1111/2041-210X.12097

Knorr, M., Frey, S., & Curtis, P. (2005). Nitrogen additions and litter decomposition: A meta analysys. Ecology, $6(12), 3252–3257. doi:10.1890/05-0150

Macdonald, E., Brummell, M. E., Bieniada, A., Elliott, J., Engering, A., Gauthier, T.-L.,… Strack, M. (2018). Using the Tea Bag Index to characterize decomposition rates in restored peatlands. Boreal Environment Research, 2469(August), 221–235.

Mori, T., Ishizuka, S., Konda, R., Wicaksono, A., Heriyanto, J., & Hardjono, A. (2015). Phosphorus addition reduced microbial respiration during the decomposition of Acacia mangium litter in South Sumatra, Indonesia. Tropics, 24(3), 113–118.

Mueller, P., Schile-Beers, L. M., Mozdzer, T. J., Chmura, G. L., Dinter, T., Kuzyakov, Y.,… Nolte, S. (2018). Global-change effects on early-stage decomposition processes in tidal wetlands-implications from a global survey using standardized litter. Biogeosciences, 15(10), 3189–3202. doi:10.5194/bg-15-3189-2018

Petraglia, A., Cacciatori, C., Chelli, S., Fenu, G., Calderisi, G., Gargano, D.,… Carbognani, M. (2019). Litter decomposition: effects of temperature driven by soil moisture and vegetation type. Plant and Soil, 435(1-2), 187–200. doi:10.1007/s11104-018-3889-x

R Core Team. (2019). R: A language and environment for statistical computing. Statistical, R Foundation for Computing, Vienna, Austria. URL https://www.R-project.org/.

Ramirez, K. S., Craine, J. M., & Fierer, N. (2012). Consistent effects of nitrogen amendments on soil microbial communities and processes across biomes. Global Change Biology, 18(6), 1918–1927. doi:10.1111/j.1365-2486.2012.02639.x

Rinkes, Z. L., Bertrand, I., Amin, B. A. Z., Grandy, A. S., Wickings, K., & Weintraub, M. N. (2016). Nitrogen alters microbial enzyme dynamics but not lignin chemistry during maize decomposition. Biogeochemistry, 128(1-2), 171–186. doi:10.1007/s10533-016-0201-0

Seelen, L. M. S., Flaim, G., Keuskamp, J., Teurlincx, S., Arias Font, R., Tolunay, D.,… de Senerpont Domis, L. N. (2019). An affordable and reliable assessment of aquatic decomposition: Tailoring the Tea Bag Index to surface waters. Water Research, 151, 31–43. doi:10.1016/j.watres.2018.11.081

Suzuki, S. N., Ataka, M., Djukic, I., Enoki, T., Fukuzawa, K., Hirota, M.,… Watanabe, K. (2019). Harmonized data on early stage litter decomposition using tea material across Japan. Ecological Research, 34(5), 575–576. doi:10.1111/1440-1703.12032

Wang, B., Blondeel, H., Baeten, L., Djukic, I., De Lombaerde, E., & Verheyen, K. (2019). Direct and understorey-mediated indirect effects of human-induced environmental changes on litter decomposition in temperate forest. Soil Biology and Biochemistry, 138(September 2018), 107579. doi:10.1016/j.soilbio.2019.107579

Wieder, R., & Lang, G. (1982). A critique of the analytical methods used in examining decomposition data obtained from litter bags. Ecology, 63(6), 1636–1642.

Xiao, W., Chen, X., Jing, X., & Zhu, B. (2018). A meta-analysis of soil extracellular enzyme activities in response to global change. Soil Biology and Biochemistry, 123(May), 21–32. doi:10.1016/j.soilbio.2018.05.001

