## Supplemental Figures for "Evaluating the tea bag method as a potential tool for detecting the effects of added nutrients and their interactions with climate on litter decomposition"

### Supporting information

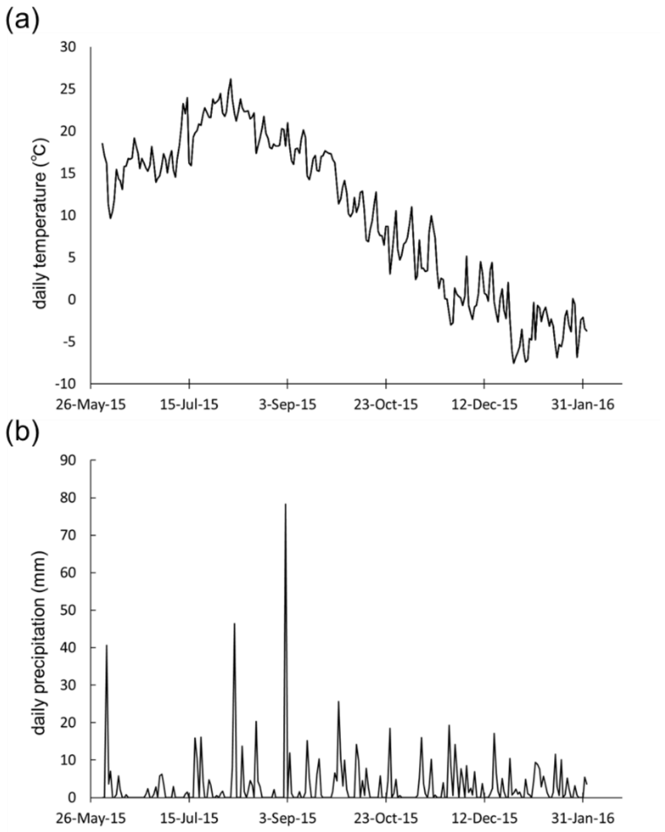


**Fig. S1.** Climate data during the experimental period.

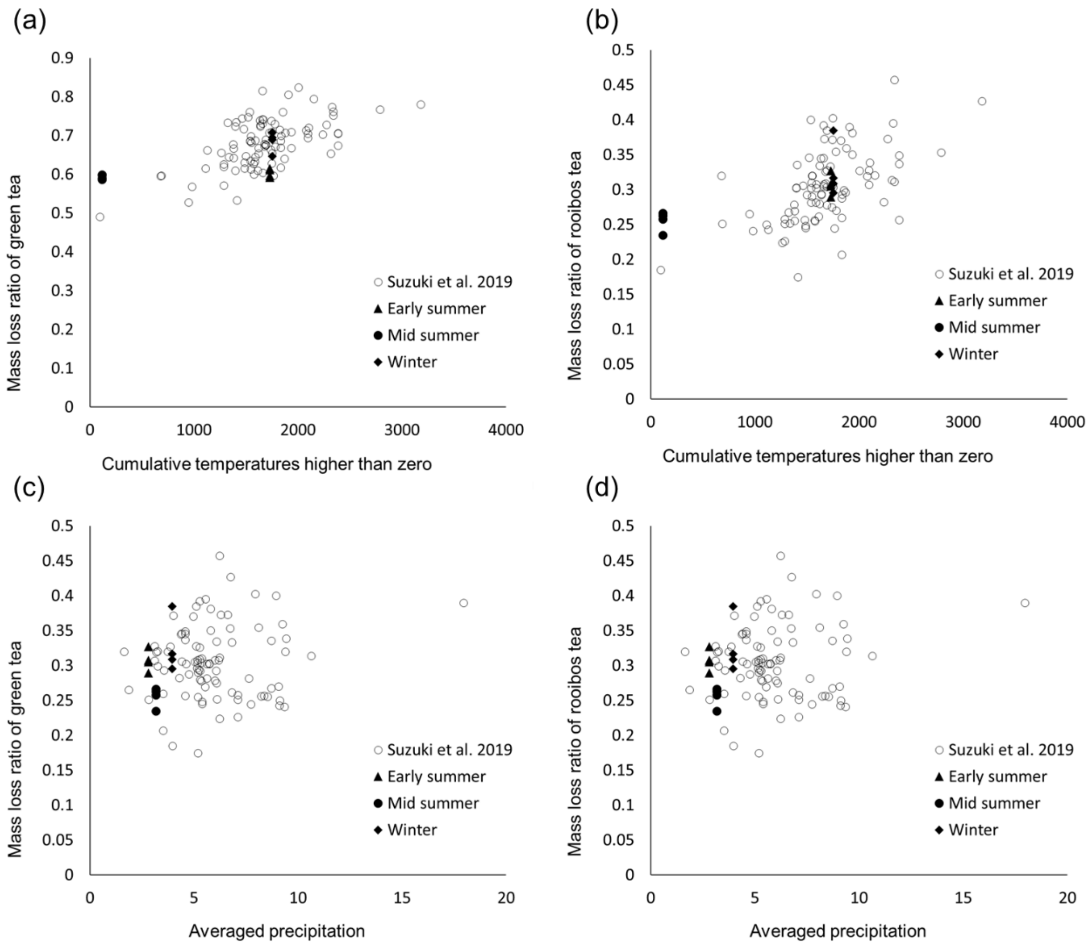


**Fig. S2.** Relationship between cumulative temperatures higher than zero or average precipitation and mass loss ratios of green or rooibos tea. Data from the present study were compared with those from other sites in Japan (Suzuki et al. 2019).

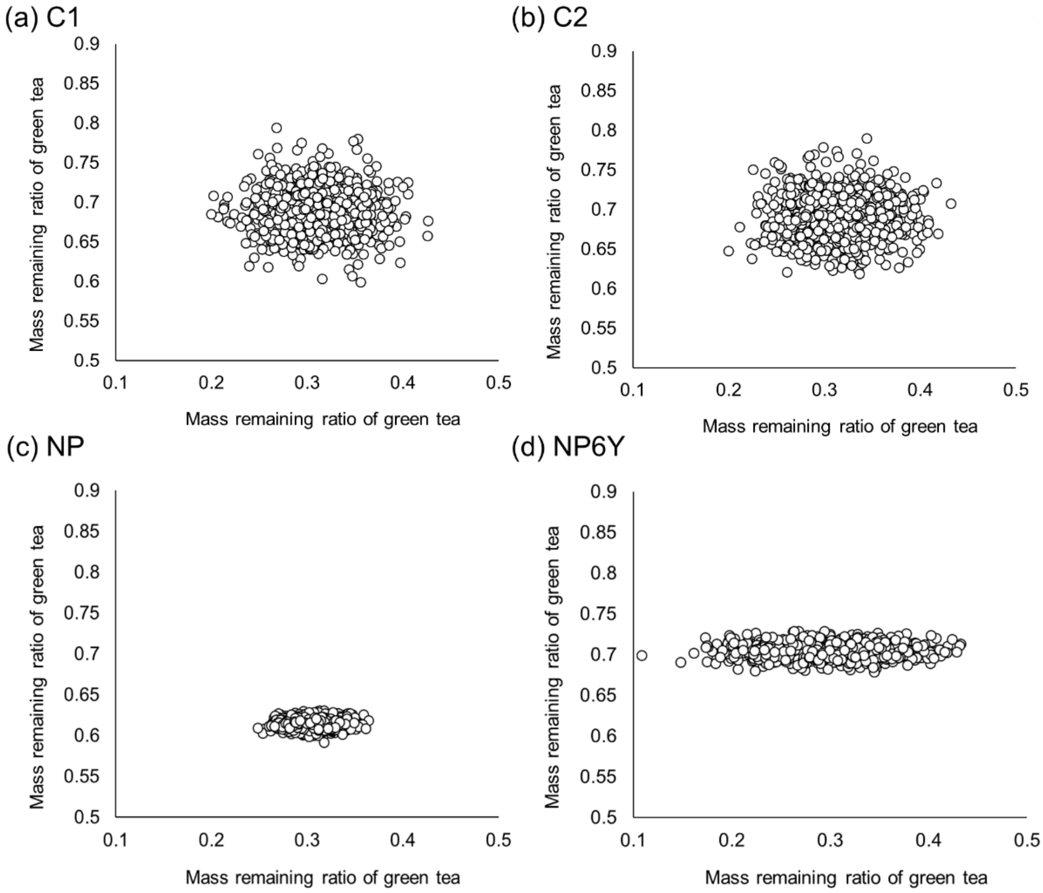


**Fig. S3.** Simulation data (n = 1000) of the ratios of green and rooibos tea mass remaining were generated using the average and standard deviation of the (a) C1, (b) C2, (c) NP, and (d) NP6Y plots in late summer, assuming a standard distribution. Each datum of the ratio of green tea mass remaining was paired with a datum of rooibos tea. We confirmed that the generated pairs did not exhibit any correlations.

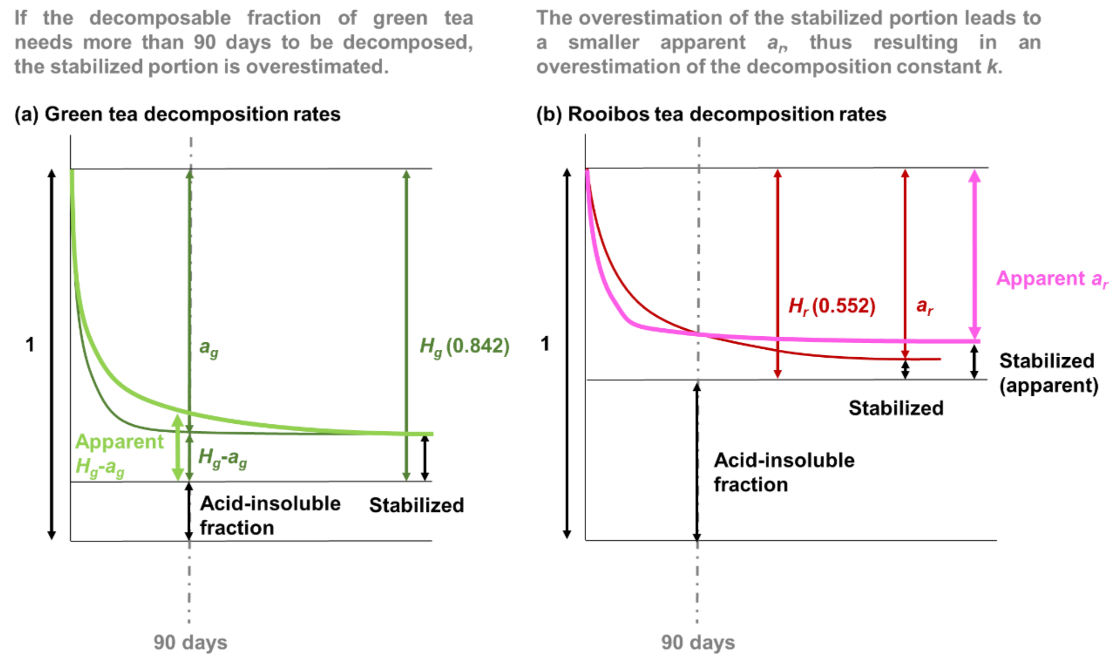


**Fig. S4.** Possible risk of a tea bag index being overestimated.
